# Obesity Human Soluble Prorenin Receptor Expressed in Adipose Tissue Improves Insulin Sensitivity and Endothelial Function in Obese Female Mice

**DOI:** 10.1101/2024.01.12.575451

**Authors:** Gertrude Arthur, Nermin Ahmed, Kellea Nichols, Audrey Poupeau, Katelyn Collins, Volkhard Lindner, Analia Loria

## Abstract

Soluble prorenin receptor (sPRR) is a component of the renin-angiotensin system (RAS) identified as a plasma biomarker for human metabolic disease. However, what tissue source of sPRR is implicated in the modulation of metabolic function remains unclear. This study investigated the contribution of human sPRR (HsPRR) produced in the adipose tissue (Adi) on the metabolic and cardiovascular function of lean and obese male and female mice. Adi-HsPRR mice, generated by crossing human sPRR-Myc-tag and Adiponectin/Cre mice, were fed a low-fat or high-fat diet (10% and 60% kCal from fat, respectively) for 20 weeks. Obese Adi-HsPRR mice showed elevated sPRR levels in adipose tissue without affecting adipocyte size or fat depot weight. Despite plasma sPRR being similar between obese Adi-HsPRR and control female mice, a positive correlation between plasma sPRR and adiposity was present only in controls. Obese Adi-HsPRR male mice showed elevated plasma sPRR compared with controls, but no correlation with adiposity was found in either group. Nevertheless, Adi-HsPRR expression improved insulin sensitivity and endothelial function, reduced adipogenic genes mRNA abundance (PPARg, SEBP1C and CD36), and increased plasma Angiotensin 1-7 levels only in obese HsPPR female mice. Taken together, elevated HsPRR in adipose tissue improved metabolic and vascular function in obese female mice despite normal circulating levels of sPRR, whereas increased local and circulating levels of HsPRR did not influence metabolic and cardiovascular function in obese male mice. Our data suggest that increased plasma sPRR associated with metabolic disease could be produced by other tissues rather than adipocytes.

## Introduction

The prevalence of obesity has reached epidemic proportions worldwide. In the United States, 2 in 5 adults are obese, accounting for 41.9% of the population (1). Obesity leads to insulin resistance and hyperinsulinemia, leptin resistance, and hyperleptinemia. Obesity is associated with a threefold higher prevalence of hypertension and is a risk factor for cardiovascular diseases (2). It is well established that obesity activates the renin-angiotensin system (RAS) through the upregulation of angiotensinogen and renin, (3–5). As such, RAS activation contributes to the development of obesity-induced diabetes, hypertension, and cardiovascular diseases (6). Prorenin receptor is a component of the RAS that is shown to promote adipogenesis and obesity (7). PRR exists in the full-length transmembrane protein, the soluble PRR (sPRR), and the truncated form (8). The full-length PRR can bind to renin and prorenin and participate in the generation of angiotensin (Ang)-I by increasing the catalytic activity of renin and promoting prorenin nonproteolytic activation (9, 10). PRR can also initiate intracellular signals, especially the activation of mitogen-activated protein (MAP) kinases through an Ang II-independent pathway (11, 12). Furthermore, PRR can interact with the membrane receptor complex of the Wnt/β-catenin pathway and with the vacuolar-type H-ATPase (13, 14).

Obesity also increases adipose tissue PRR expression (15, 16) in humans; however, studies in mice showed that the functional role of PRR in adipogenesis includes Ang II generation, extracellular signal-regulated kinase (ERK1/2) activation, and regulation of the peroxisome proliferator-activated receptor gamma (PPARᵧ), known as the master regulator of adipogenesis (17, 18). The loss of PRR in adipose tissue reduces peroxisome proliferator activated receptor gamma (PPARᵧ) expression (17) and PPARᵧ knockdown decreases adipose PRR expression (19).

Ablation of the PRR gene has been shown to attenuate metabolic disease. In the pancreas, the knockdown of PRR in β-cells resulted in diabetes by reducing insulin secretion and impairing glucose sensitivity (20). In the adipose tissue, PRR knockdown improved insulin sensitivity in male mice on a normal diet and female mice on a high fat diet (16). Together, these studies show that PRR expression is required for normal adipogenesis in physiological conditions, while increased PRR expression promotes glucose intolerance and insulin resistance.

In humans, elevated circulating sPRR is present in obese individuals (21), whereas weight loss after bariatric surgery has been shown to reduce its circulating levels (21). Additionally, plasma sPRR is elevated in women with type 2 diabetes (22), as well as in hypertensive patients on RAS blockers, suggesting that elevated circulating sPRR in humans is associated with the effects of obesity on blood pressure regulation. However, the influence of sPRR on obesity, glucose, and insulin homeostasis is still under debate. Several studies have shown that increased circulating sPRR in obesity settings induces glucose intolerance and insulin resistance. Accordingly, whole-body loss of sPRR reduced body weight (23), while a chronic sPRR infusion abolished this effect (24). Furthermore, HFD-induced obesity elevated plasma sPRR leading to hyperglycemia, hyperinsulinemia, and glucose intolerance, characteristic of type 2 diabetes (T2D) (25). Conversely, others have shown that sPRR infusion in a model HFD-induced obesity, reduced body weight and improved glucose and insulin sensitivity by increasing adipose glucose transporter type 4 (Glut 4) and the phosphorylation of proteins kinase B (Akt) (19). Infusion of sPRR in high fat diet-induced obesity also increased blood pressure (BP) (24, 26) and induced endothelial dysfunction via the activation of the angiotensin II type 1 (AT1) receptor (26).

Strikingly, all the research studies mentioned above that tested the impact of sPPR were conducted only on male mice using whole-body approaches. To address this gap, we developed a mouse model that specifically expresses human sPRR (HsPRR) in adipose tissue (Adi-HsPRR). We used both male and female mice expressing Adi-HsPRR and subjected them to low or high-fat diets for 20 weeks to investigate the effects on adipose tissue remodeling, glucose and insulin homeostasis, endothelial function, and blood pressure.

## Methods

The original data that support the findings of this study are available from the corresponding author upon reasonable request.

### Transgenic mouse model

All animal studies described below complied with the University of Kentucky Institutional Animal Care and Use Committee (Protocol No. 2022-4059) and followed the “Guide for the Care and Use of Laboratory Animals” issued by the National Institutes of Health.

Mice were on 14:10 hours, light: dark cycle, starting at 7:00 am for the light cycle and at 9:00 pm for the dark cycle. Mice had *ad libitum* access to water and diet.

Human sPRR-Myc-tag transgenic mice were developed by cloning myc epitope-tagged sPRR into the CAG-GFP vector (27). The fragment was sequenced to the confirm accuracy of the construct and transfected into HEK293 cells stably expressing Cre recombinase and screened with a myc monoclonal antibody (clone Vli01, MaineHealth Institute for Research, Scarborough, ME) to verify the expression of the fragment in the media. Upon validation, the construct was injected into oocytes of C57BL/6J mice to generate male and female Human sPRR-Myc-tagged transgenic mice.

Adi-HsPRR male and female mice were generated by breeding heterozygote female human sPRR-Myc-tag transgenic mice with male mice expressing Cre recombinase under the control of the Adiponectin promoter (B6; FVB-Tg (AdipoCre)1Evdr/J, Jax No. 010803, The Jackson Laboratory, Figure 1A).

**Figure 1.**
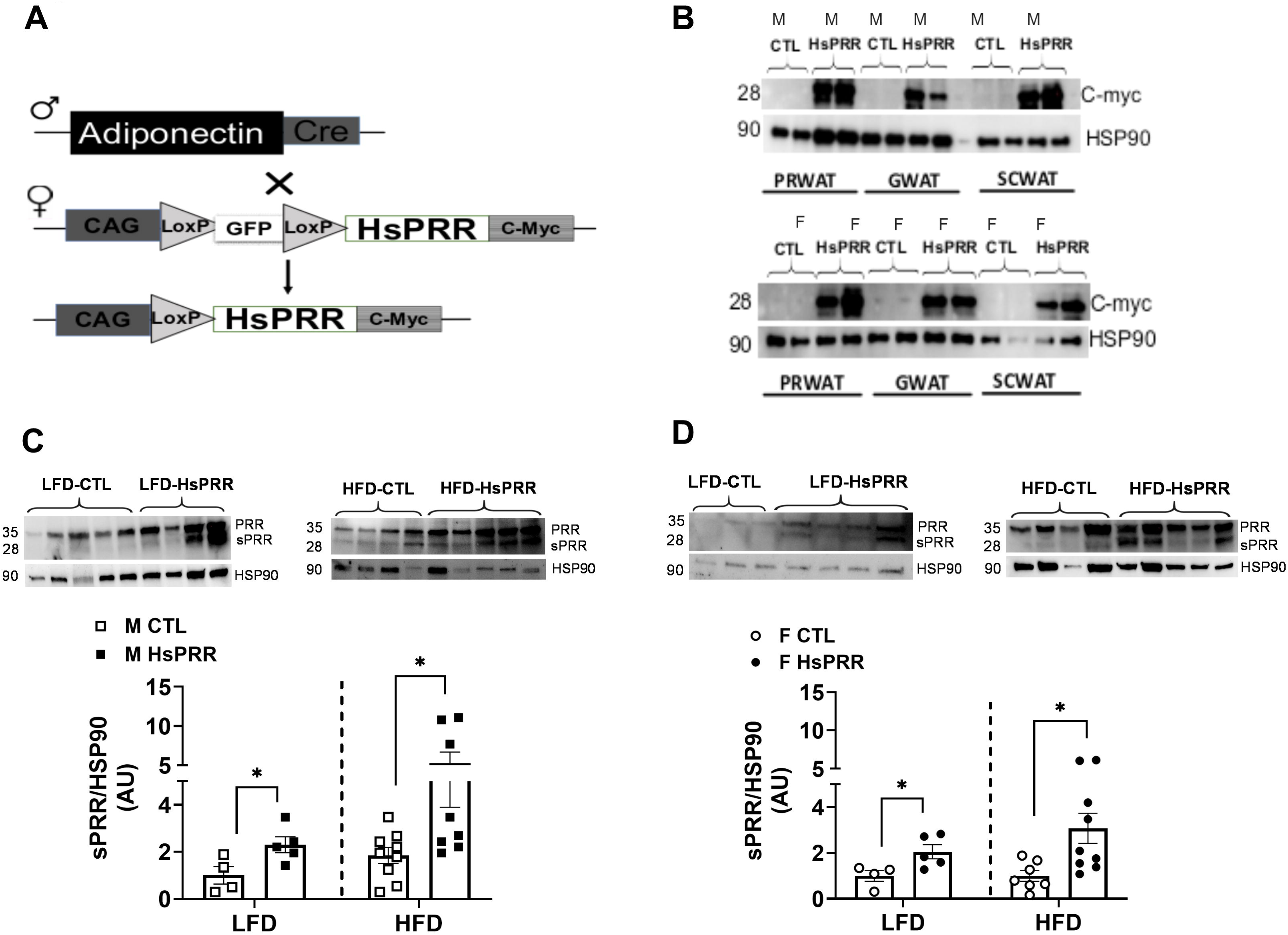
Generation and validation of adipose-specific human sPRR (Adi-HsPRR) mouse model. (A) Adipose tissue-derived human sPRR male and female mice were generated by breeding heterozygote male mice that expressed Cre recombinase under the control of the Adiponectin promoter with female heterozygote mice that expressed the human soluble prorenin receptor (HsPRR/+). (B) Expression of Human sPRR was validated by the detection of myc-tagged HsPRR in adipose tissue (peri-renal, gonadal, and subcutaneous). Tissue sPRR expression was assessed via western blot in (C) Male and (D) Female mice. Data are expressed as mean ± SEM of 7-1 mice/group. A 2-way ANOVA was performed to detect differences

### Experimental Design

Adi-HsPRR and control littermates (CTL) male and female mice (Males= 9-15/group; Females=8-14/group) were weaned and kept on a regular chow diet for 8 weeks (18% protein, Teklad Global Diet, ENVIGO Harlan, Madison, WI). Then, mice were switched to either a low-fat diet (10% kcal from fat; D12450J, Research Diets, Inc. USA) or a high-fat diet (60% kcal from fat; D12492, Research Diets, Inc. USA) for 20 weeks. Body weight was recorded weekly and body composition (EchoMRI) was assessed monthly. The intraperitoneal glucose tolerance test (ipGTT) was performed at 12 weeks on special diets and the insulin tolerance test (ITT) was performed at 14 weeks. At 16 weeks, mice were housed individually and implanted in the left carotid artery with a catheter connected to a telemetry transmitter (PA-C10 model; DSI; Saint Paul, MN). Surgeries were performed under isoflurane anesthesia and mice were treated with meloxicam for 2 days (4mg/kg daily, subcutaneous injection). One week after recovery, systolic blood pressure (SBP), mean arterial pressure (MAP), diastolic blood pressure (DBP), and heart rate (HR) were measured for 5 consecutive days using the DSI Ponemah software. At the end of the study, tissues and plasma were collected for biochemical analysis and vascular studies.

### Intraperitoneal Glucose and Insulin Tolerance Test

Intraperitoneal glucose tolerance test (ipGTT) was performed on mice after a 6-h fast (28) and intraperitoneal insulin tolerance test (ITT) was performed after a 3-h fast (29). Blood glucose concentration was measured at baseline using a glucometer (Accu-Chek Aviva Plus, Roche Diagnostics). Following glucose (1g/kg, i.p) and insulin injection (i.p), blood glucose was assessed at 15-, 30-, 60- and 120 minutes. Insulin dose is as follows, LFD males: 0.75 IU/kg; LFD females: 0.4 IU/kg; HFD males: 1 IU/kg; HFD females: 0.9 IU/kg.

### Vascular Reactivity

Vascular reactivity of the thoracic aorta was measured as previously reported (30). Mouse thoracic aorta was isolated, and cleaned from perivascular adipose tissue. Rings were cut into 4 concentric rings (∼3 mm each), mounted on pins for wire myography (Danish Myo Technology A/S, Aarhus, Denmark), and tested for viability; preconstricted with serotonin (SE, 1×10^-4^ M) and relaxed with acetylcholine (ACh, 1×10^-3^ M). Cumulative concentration-response (CCR) curves to potassium chloride (KCl, 4.7 mM to 100 mM) were utilized to assess constriction. Endothelial-dependent relaxation was assessed, aortic segments were pre-constricted with SE (1x10^-4^ M, 5 ul), and CCR curves to ACh (1×10^-9^ to 1×10^-5^ M) were performed. Following recovery, vasorelaxation to the endothelium-independent vasodilator sodium nitroprusside (SNP; 1×10^−10^ M to 1×10^-5^ M) was assessed in the same aortic segments. Vasorelaxation data are presented as percent relaxation from SE-induced constriction. For all CCR curves, maximum response (Emax) was calculated using Prism Software (GraphPad, San Diego, CA).

### Immunostaining

Immunostaining was performed on gonadal white adipose tissue (gWAT) from male and female HFD mice as previously reported (31). Gonadal WAT was fixed in paraformaldehyde and embedded in paraffin blocks. Sections were stained with hematoxylin and eosin and examined with a Nikon Eclipse 80i light microscope. All sections were photographed and quantified using NIS Elements BR.3.10 software to determine cell size and count.

### Quantification of Plasma Parameters

Plasma sPRR was determined by soluble (Pro)renin Receptor ELISA (IBL, Minneapolis, MN) as previously reported (32). Plasma renin was determined by Mouse Renin ELISA (IBL, Minneapolis, MN) (32). Plasma RAS levels were assessed using LC-MS/MS at the University of Kentucky RAAS Analytical Laboratory as previously described (33).

### Immunoblotting

Protein from frozen retroperitoneal adipose tissue was extracted and 40µg of protein was run on precast polyacrylamide gel as previously described (31) and then transferred to a polyvinylidene difluoride membrane via a semi-wet transfer equipment (Trans-Blot Turbo Transfer System; Bio-Rad Laboratories, Hercules, CA) as recommended by the manufacturer. Membranes were blocked in 5% nonfat dried milk in Tris-buffered saline with 0.1% Tween 20, membranes were incubated with C-myc antibody (MaineHealth Institute for Research, clone Vli01) and HSP90 (Cell signaling, #4874) in 5% nonfat dried milk in Tris-buffered saline with 0.1% Tween 20. After incubation with handle region decoy peptide-conjugated antirabbit secondary antibody (Jackson ImmunoResearch Laboratories, AB2313567), proteins were imaged using the ChemiDoc Imaging System (Bio-Rad Laboratories, Hercules, CA). The levels of proteins were quantified using Image Lab software (Bio-Rad Laboratories, Hercules, CA) and normalized to HSP90.

### Tissue RNA Extraction and Quantitative RT-PCR

RNA from gonadal white adipose tissue was extracted with the SV Total RNA Isolation System (Promega, Madison, WI) as previously reported (34). RNA concentration was measured by a NanoDrop (Thermo-Fisher Scientific, Madison, WI), and cDNA was prepared from 500ng of RNA using Qscript cDNA SuperMix (Quanta Biosciences, Gaithersburg, MD). mRNA expression of genes was evaluated by quantitative RT-PCR using PerfeCTa SYBR Green FastMix (Quanta Biosciences) and normalized to 18S.

### Statistical Analysis

Data are represented as means±SEM. Statistical analysis was performed using Graph Prism (GraphPad Prism 10.2; San Diego, CA, USA). Statistical differences between groups were assessed by a two-way ANOVA followed by Tukey post hoc analysis for multiple comparisons. Grubbs test (GraphPad Quick-Calcs) was used to determine statistical outliers. Values of P<0.05 were considered statistically significant.

## Results

### Characterization of Adi-HsPRR mouse model

Figure 1A shows the breeding strategy to generate Adi-HsPRR mice. To confirm the expression of human myc-tagged sPRR in the adipose tissue of mice, a western blot analysis of c-myc was performed. The protein expression of myc-tagged HsPRR was detected at 28 KDa in perirenal, gonadal, and subcutaneous adipose tissue of male and female Adi-HsPRR mice (Figure 1B) and was absent in control male and female littermates. Expression of sPRR in perirenal adipose tissue was increased in male and female Adi-HsPRR mice on low- and high-fat diets (Figure 1C-D). sPRR expression was also increased in subcutaneous and gonadal adipose tissue (Supplementary Figure 1).

### Adi-HsPRR expression does not influence the morphology and size of the adipocytes

The body weight, fat, and lean mass of both CTL and Adi-HsPRR mice were similar in both lean and obese mice (as mentioned in Supplementary Table 1). Additionally, the weights (Figure 2A-B), adipocyte size (Figure 2C-D), and morphology (Supplementary Figure 2) of perigonadal white adipose tissue (gWAT) were also similar between control and Adi-HsPRR mice. As a result, plasma leptin levels were also found to be similar between the two groups (Supplementary Figure 3A-B).

**Figure 2.**
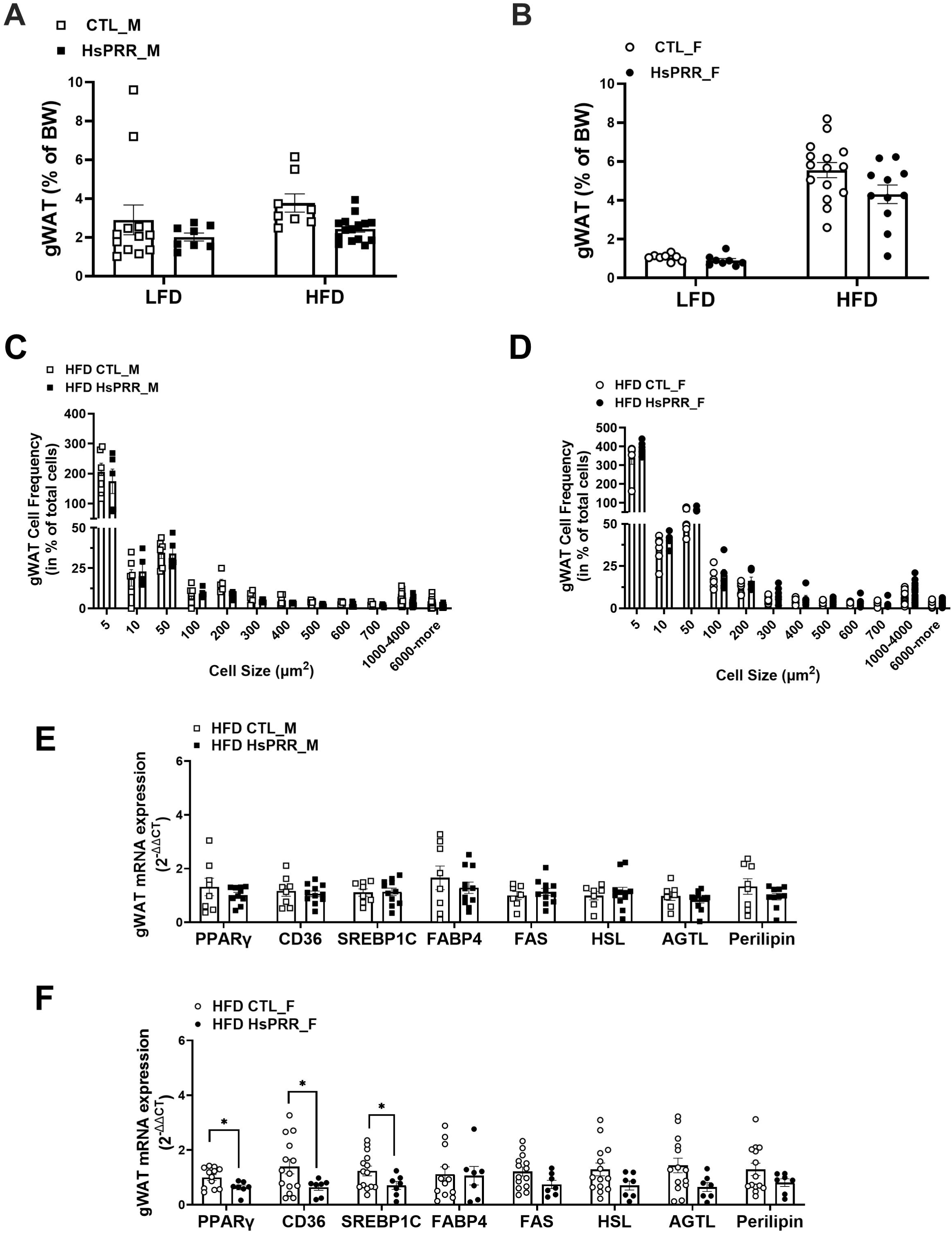
Effect of Adi-HsPRR expression of Adipose Tissue. Fat pads at takedown in (A) Male and (B) Female mice. Data are expressed as mean ± SEM of 8-16 mice/group. A 3-way ANOVA was performed to detect differences. Tissue morphology of gonadal white adipose tissue was analyzed and quantified in (C) Male HFD and (D) Female HFD mice using Nikon NIS Elements BR.3.10 software at 10X (200µm). Data are expressed as mean ± SEM of 5-15 mice/group. Gene expression analysis was performed in gonadal WAT of (E) Male HFD and (F) Female HFD mice. Data are expressed as mean ± SEM of 8-15 mice/group. A t-test was performed to detect differences.

In obese male mice, the mRNA expression of various transcription factors and genes involved in adipogenesis, lipid metabolism, fatty acid synthase (FAS), hormone-sensitive lipase (HSL), adipose triglyceride lipase (ATGL), and perilipin were similar to those in the same fat depot. However, in obese female mice expressing HsPRR, scavenger receptor class B member 3 (CD36) and sterol regulatory element-binding transcription factor 1C (SREBP1C) were reduced in gWAT compared to the male group. This was noted in Figure 2E-F of the study.

### Adi-HsPRR modulates circulating RAS in obese mice

Although mice fed a low-fat diet showed similar levels between genotypes, a high-fat diet increased sPRR levels in both control male mice. However, plasma sPRR was further increased in Adi-HsPRR mice as depicted in Figure 3A. In females, HFD induced a similar increase in plasma sPRR between genotypes as shown in Figure 3B. Plasma renin concentration (PRC) was reduced similarly between genotype and sex in response to HFD, as demonstrated in Figure 3C-D. However, the expression of Adi-HsPRR reduced Ang 1-7 levels in obese male mice and increased it in obese female mice, as illustrated in Figure 3D-E. Other angiotensin peptides in mice fed a HFD were similar between groups, including plasma renin activity, plasma aldosterone, and angiotensin peptides: Ang II (1–8), Ang I (1–10), Ang III (2–8), Ang 1-5, Ang IV (3–8), Ang 1-9, Ang 2-10, which were found to be similar between HFD groups as per Supplementary figure 4C-D.

**Figure 3.**
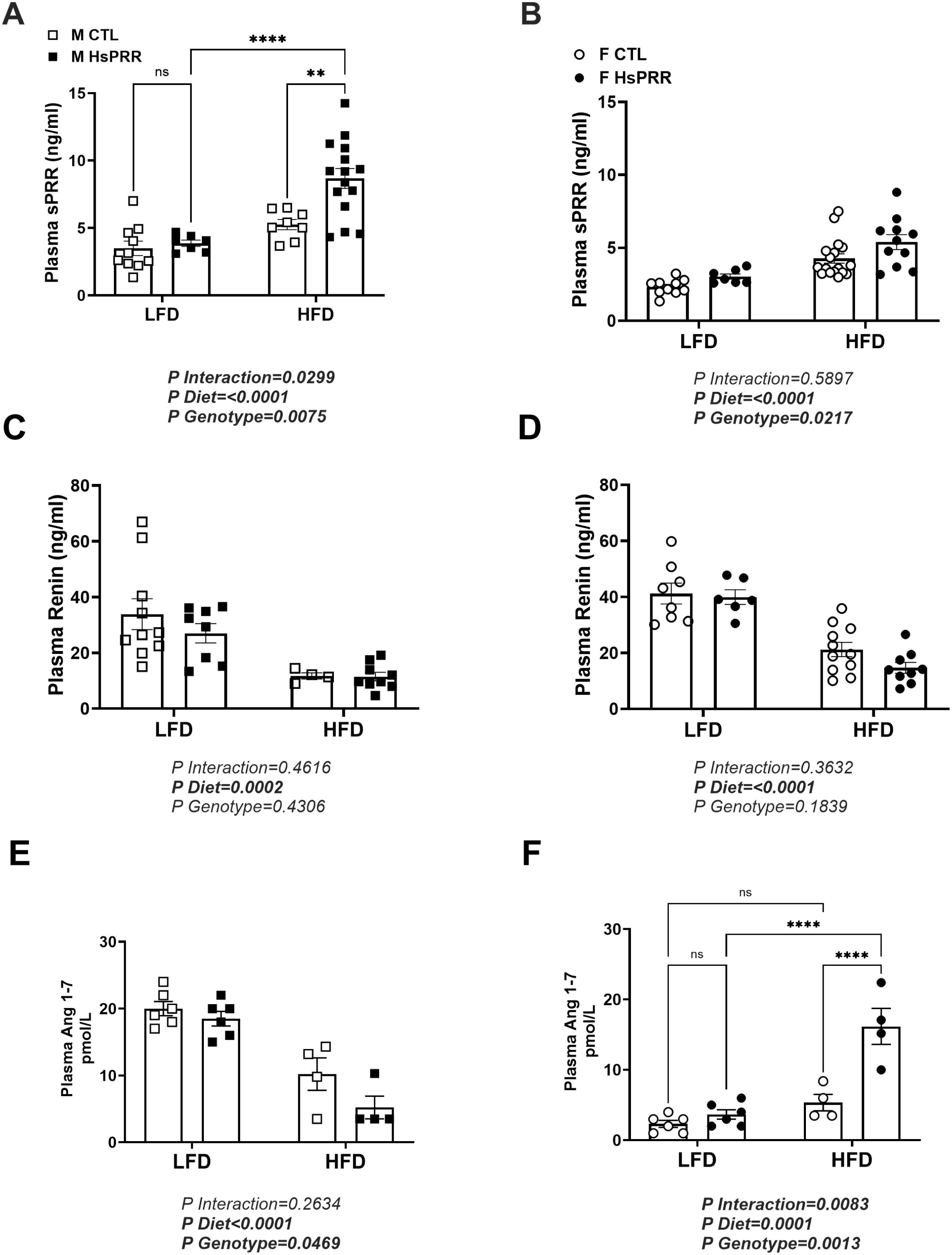
Adi-HsPRR expression modulates circulating RAS. Plasma sPRR in (A) Male and (B) Female mice. Data are expressed as mean ± SEM of 7-17 mice/group. Plasma renin in (C) Male and (D) Female mice. Data are expressed as mean ± SEM of 5-11 mice/group. Plasma Ang 1-7 in (E) Male and (F) Female mice. Data are expressed as mean ± SEM of 4 mice/group. A 2-way ANOVA was performed to detect differences

### Increased adiposity in obese females correlates with circulating sPRR

Male mice did not show any correlation between visceral adiposity (gWAT, %BW) and changes in plasma sPRR levels, regardless of their diet or genotype (Figure 4A). In female CTL and Adi-HsPRR mice fed with a LFD, there was no association between gWAT and plasma sPRR as well. However, female CTL mice fed a HFD showed a positive correlation between visceral adiposity and plasma sPRR, which was voided in obese Adi-HsPRR female mice (Figure 4B).

**Figure 4.**
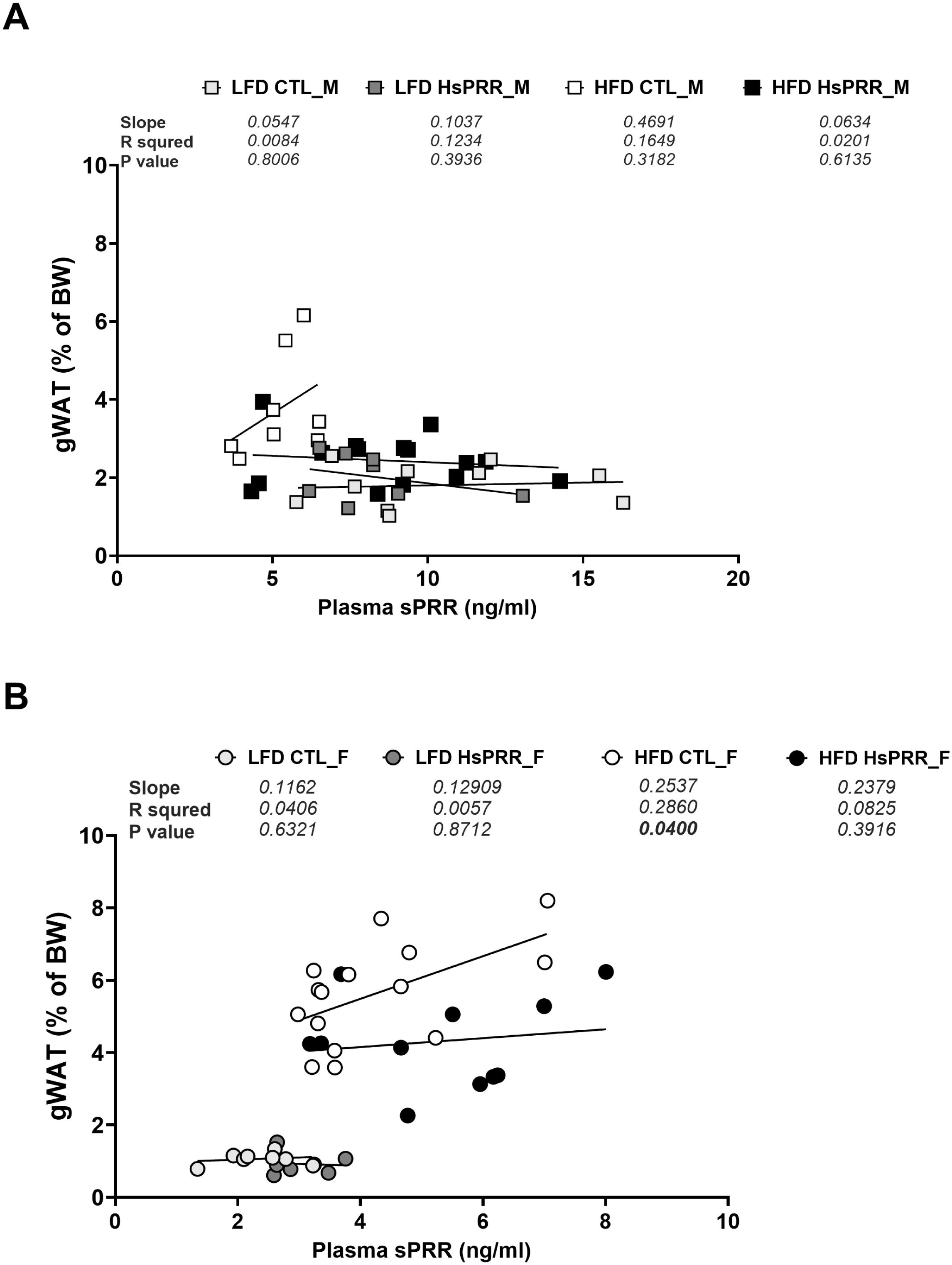
Increased plasma sPRR correlates with adiposity in obese females. Linear regression of visceral adiposity (gWAT) and plasma sPRR in (A) Male mice, and (B) Female mice. A simple linear regression was performed to detect differences.

### Adi-HsPRR improves insulin sensitivity in female mice fed a HFD

Fasting blood glucose levels were similar between CTL and Adi-HsPRR mice in both males and females (as shown in Supplementary Figure 5A-D), and HFD increased blood glucose further in all groups. Moreover, the time-dependent glucose clearance remained similar between the two groups, whether they were on a low-fat or high-fat diet (as illustrated in Figure 5A-B). Interestingly, glucose utilization upon insulin challenge was found to be comparable in obese male mice regardless of HsPRR expression in adipose tissue (as demonstrated in Figure 5C). However, the expression of HsPRR in adipose tissue led to an improvement in blood glucose utilization in obese female mice compared to CLT littermates (as depicted in Figure 5D). This suggests that obese Adi-HsPRR female mice have better insulin sensitivity.

**Figure 5.**
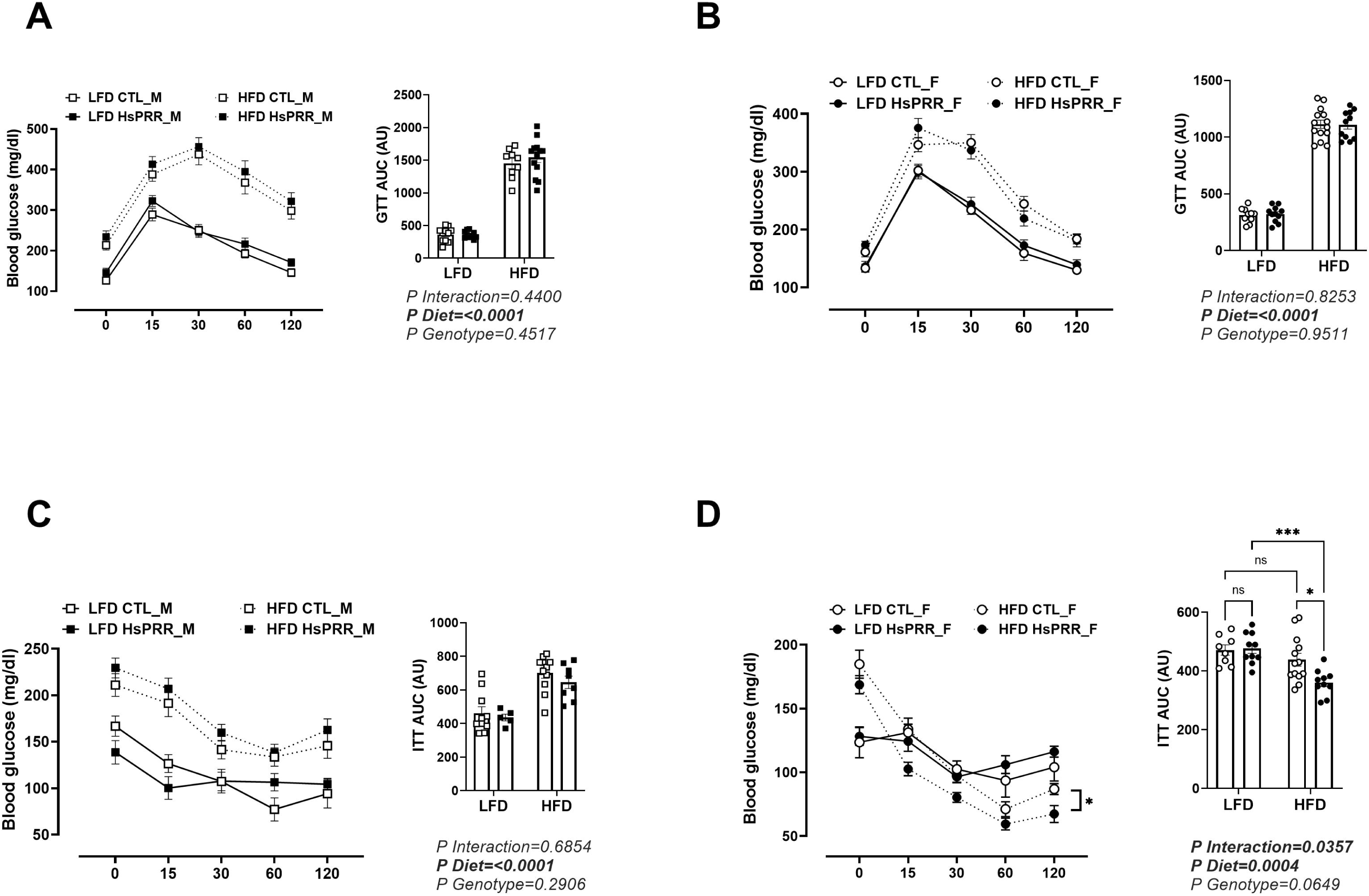
Adi-HsPRR improves insulin sensitivity in HFD females. Time-dependent glycemia and area under the curve after glucose injection in (A) Male and (B) Female mice. Data are expressed as mean ± SEM of 9-14 mice/group. Time-dependent glycemia and area under the curve after insulin injection in (C) Male and (D) Female mice. Data are expressed as mean ± SEM of 6-13 mice/group. A 2-way ANOVA was performed to detect differences.

### Adi-HsPRR improves endothelial function in female Adi-HsPRR mice fed a HFD

Endothelium-dependent relaxation of the thoracic aorta with acetylcholine (ACh) was similarly impaired in obese male CTL and Adi-HsPRR mice (Figure 6A). However, obese female Adi-HsPRR counterparts displayed better endothelium-dependent vascular relaxation compared to CTLs (Figure 6B). Endothelium-independent relaxation with sodium nitroprusside (SNP) was similar between genotypes in male and female mice (Figure 6A-B). In addition, maximal potassium chloride (KCl) was similar between groups in males and females (Supplementary Figure 5A-B).

**Figure 6.**
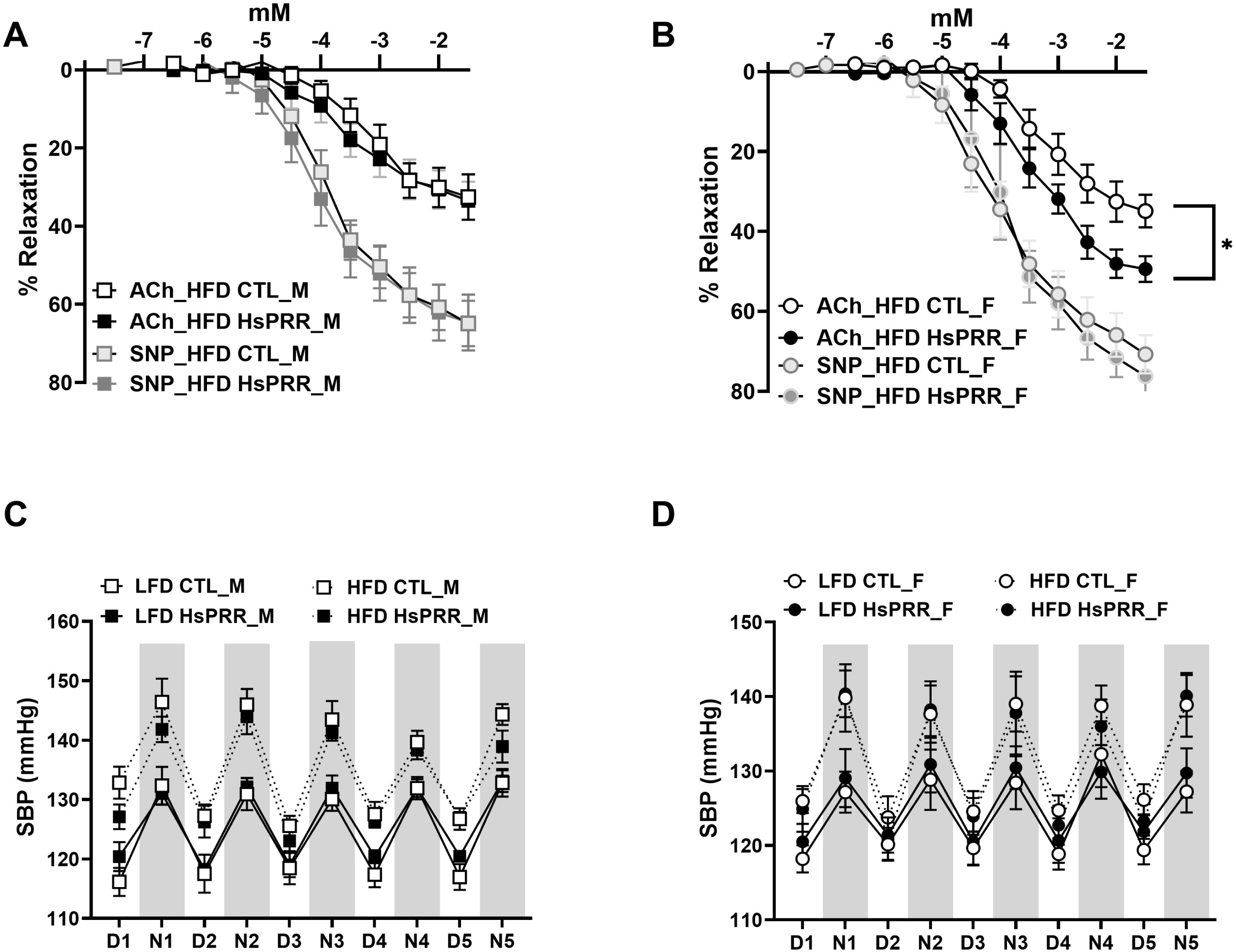
Adi-HsPRR improves endothelium-dependent vascular relaxation in HFD females. Vascular relaxation in (A) Male and (B) Female mice. Data are expressed as mean ± SEM of 5 mice/group. A 2-way repeat ANOVA was performed to detect differences. Day and night systolic blood pressure in (A) Male and (B) Female mice. Data are expressed as mean ± SEM of 4-9 mice/group. A 3-way repeat ANOVA was performed to detect differences.

Obesity-induced hypertension was present in both male and female mice regardless of the genotype (Figure 6B-C) and it was not influenced by HsPRR expression in adipose tissue.

## Discussion

In this study, we report for first time that adipose HsPRR expression increased tissue sPRR expression, but did not have any effect on body weight, adipose tissue morphology, or plasma leptin in mice on low or high-fat diets. In addition, circulating sPRR levels were similar between groups fed a low-fat diet; however, we observed that a HFD increased circulating sPRR levels in all groups. Although Adi-HsPRR further increased plasma sPRR levels only in HFD male mice, only obese female control mice showed a correlation between body adiposity and plasma sPRR, indicating that increased circulating sPRR may not be the primary cause of increased adiposity during obesity in males, but can play a role in fat expansion during the development of obesity in females. Moreover, it suggests that the influence of circulating HsPRR may be negligible compared to its local effect. This is in line with our recent report that local expression of HsPRR in the kidney exerts a cardio-renal effect independent of circulating sPRR (32). Furthermore, obese Adi-HsPRR female mice displayed insulin sensitivity and vascular relaxation, increased levels of the vasodilator agent, Ang 1-7, and reduced CD36 and SREBP1C gene expression in gonadal white adipose. Overall, these findings suggest that HsPRR in adipose tissue exerts a sex-specific effect on insulin sensitivity and endothelial function, primarily through local rather than systemic effects of HsPRR.

The adipose tissue and liver have been previously demonstrated as important sources of circulating sPRR (31, 34). In clinical studies, plasma sPRR is elevated in obese subjects but decreases after weight loss due to bariatric surgery (21, 35). Likewise, in our study, obese male and female mice displayed increased plasma sPRR compared to their lean controls. This data suggests that adipose tissue-derived-HsPRR is an important contributor to the increased sPRR circulating in obese mice, but adipose tissue-derived sPRR seem unlikely to play a role in metabolic dysfunction. Thus, whether metabolic dysfunction during obesity is impaired when sPRR production is increased in other tissues needs further investigation.

In obese male mice, infusion of sPRR was shown to reduce gonadal white adipose tissue (19). Our study shows that the expression of human sPRR in adipose tissue did not result in any significant change in body weight or composition between CTL and Adi-HsPRR mice. Moreover, it did not affect the morphology of adipose tissue, despite the fact that plasma sPRR was robustly elevated in obese male Adi-HsPRR mice. Reasons for this discrepancy could be the length of HFD feeding (20 weeks in our study vs 32 weeks used in other studies), method of elevated plasma sPRR generation (increased tissue derived vs infusion), or species differences (mouse vs human sPRR). Additionally, other studies have demonstrated that other factors such as PRR and PPARᵧ may play a bigger role in adipogenesis and obesity than sPRR. Indeed, in a lipodystrophy model induced by knockdown of PRR in adipose tissue, plasma sPRR was elevated (17). Moreover, infusion of sPRR in PPARᵧ KO mice, another model of lipodystrophy, did not affect body weight (19).

Clinical studies have shown that plasma sPRR is elevated in women but not men with type 2 diabetes (22). Moreover, increased circulating sPRR is associated with gestational diabetes (36). However, these studies do not show an association between plasma sPRR and glucose metabolism in men (22). Conversely, obese male mice with hyperglycemia and hyperinsulinemia display elevated plasma sPRR (25), while the exogenous addition of sPRR promote adipose tissue beigeing (37) and improve glucose and insulin balance in morbidly obese male mice (19). In the present study, we found that human sPRR expression in adipose tissue did not affect glucose and insulin metabolism irrespective of circulating sPRR levels, suggesting that plasma human sPRR in males may not play a role in glucose and insulin homeostasis, similar to reported findings in human studies (22). The discrepancies in findings between human and mouse sPRR may be due to species differences that need further investigation. Although the sPRR gene has more than 90% identity between humans and mice, slight differences could lead to variations in signaling and response.

Adi-HsPRR also reduced SREBP1c and CD36 gene expression and increased vasodilator agent Ang 1-7 in obese female mice. Ang 1-7, produced from the cleavage of Ang II by ACE2 and the cleavage of Ang I by neprilysin (NEP) upon binding to the Mas receptor induces vasodilatation, antihypertrophic, antiinflammation, anti-remodeling, and antiapoptotic effects (38). Recent evidence shows that Ang 1-7 improves metabolic function and its levels could be a marker of metabolic syndrome (39). Increased Ang 1-7 levels have been shown to improve skeletal muscle insulin sensitivity (40) and protect against streptozotocin-induced diabetes mellitus in rats through improvements in pancreatic β cell survival, insulin secretion, and utilization (41). Moreover, as a potent vasodilator, antihypertrophic, antiinflammation, and anti-remodeling agent, Ang 1-7 has been shown to improve endothelial function. Ang 1-7 decreases Ang II-induced matrix metalloproteinase 8 (MMP-8) expression which drives atherosclerosis (42). In humans, an imbalance between Ang II and Ang 1-7 has been associated with endothelial dysfunction and inflammation in diabetics (43). Hence, the increase in circulating Ang 1- 7 in female Adi-HsPRR mice could be a primary pathway or mechanism for improved insulin response and endothelial function in Adi-HsPRR female mice. Notably, male Adi-HsPRR mice, which show decreased Ang 1-7 levels, do not have an improvement in insulin response and endothelial function. Upcoming studies will investigate the mechanism of HsPRR-Ang 1-7 mediated insulin signaling and vascular functioning.

The role of SREBP1c in lipid metabolism and insulin signaling is well documented (44, 45). Increased expression of SREBP1c in the adipose tissue of mice has been shown to lead to insulin resistance and type 2 diabetes (46). In humans, hyperinsulinemia increased SREBP1c gene expression in white adipose tissue (47). Likewise, our findings show that increased insulin sensitivity, induced by HsPRR expression in the adipose tissue decreased SREBP1c expression. SREBP1c is downstream of insulin signaling, hence its presence could be stimulated by either the insulin receptor-Akt or the insulin receptor-PKCλ pathway (48). The exact mechanism of SREBP1c reduction in Adi-HsPRR female mice would be the focus of future studies. CD36 is an important factor in fatty acid uptake. Increased expression of CD36 in the adipose tissue has been linked to insulin resistance in *vitro* (*49*) and humans (50, 51). Similarly, a decrease in CD36 expression in gonadal WAT of Adi-HsPRR females is linked to improved insulin responses observed.

Our study has revealed important insights into the intricate interplay between adipose-derived human sPRR, circulating RAS, and lipid trafficking regulators in the adipose tissue. We have identified that this interaction has a direct effect on insulin signaling and vascular function, and that these effects are sex-specific. Our results suggest that the model we developed could be a novel tool for further investigating the potential antidiabetic effects of adipose sPRR in obese women. Our findings also indicates that sPRR, which is a protein that regulates the RAS, plays a role in the development of insulin resistance and vascular dysfunction in obese individuals. By studying this potential mechanism further, we may be able to develop new treatments for diabetes and other related metabolic disorders. Overall, our findings have important implications for the management of obesity-related metabolic health, particularly in women.

## Acknowledgments

We thank the Mouse Genome Modification Core (supported by COBRE 1P20GM121301, L. Liaw PI) at the MaineHealth Institute for Research for the generation of the Cre-inducible human sPRR-myc-tag transgenic mouse strain. We thank the UK RAAS Analytical Laboratory for assistance with plasma RAS measurement.

## Disclaimers and Disclosures

None

## Supplemental Materials

Tables 1 S1-S6

