## Supplementary data for "Obesity Human Soluble Prorenin Receptor Expressed in Adipose Tissue Improves Insulin Sensitivity and Endothelial Function in Obese Female Mice"

### Figures and Figure Legends

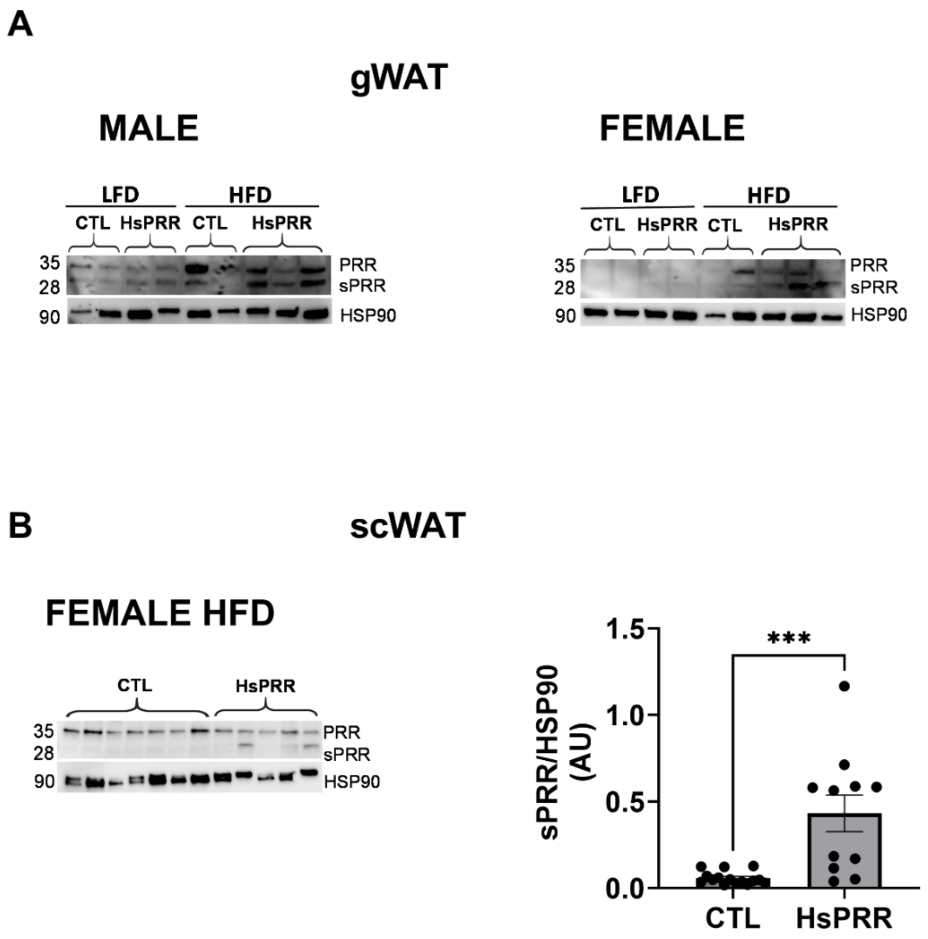


**Supplementary Figure 1 Expression of sPRR in adipose tissue**

sPRR expression in (A) gonadal and (B) subcutaneous white adipose tissue in male and female mice.

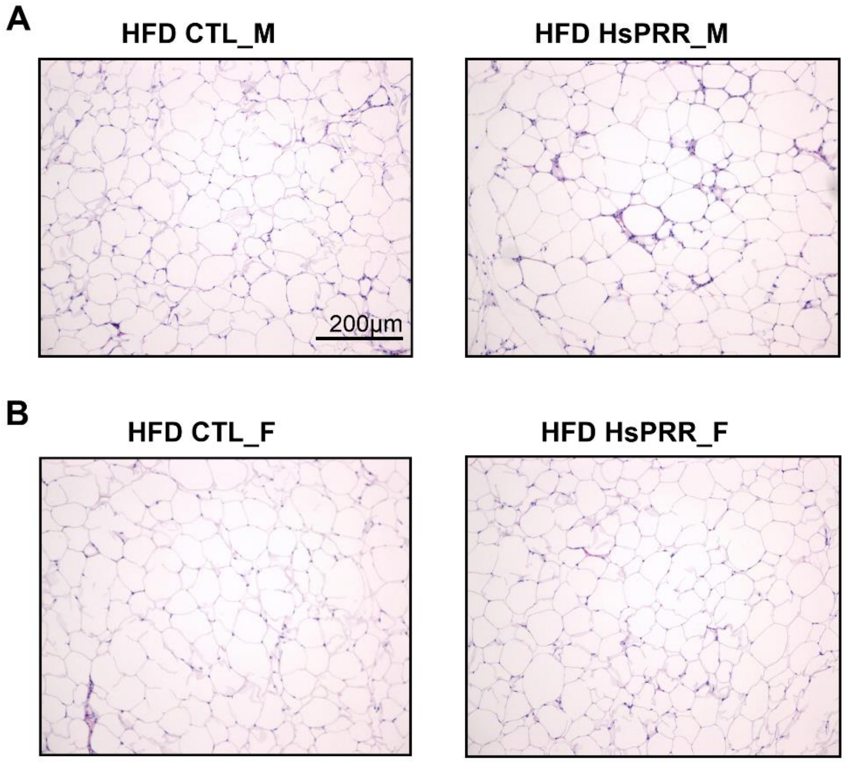


**Supplementary Figure 2 Histology of gonadal WAT from obese mice**

Histology from (A) Male HFD and (B) Female HFD mice.

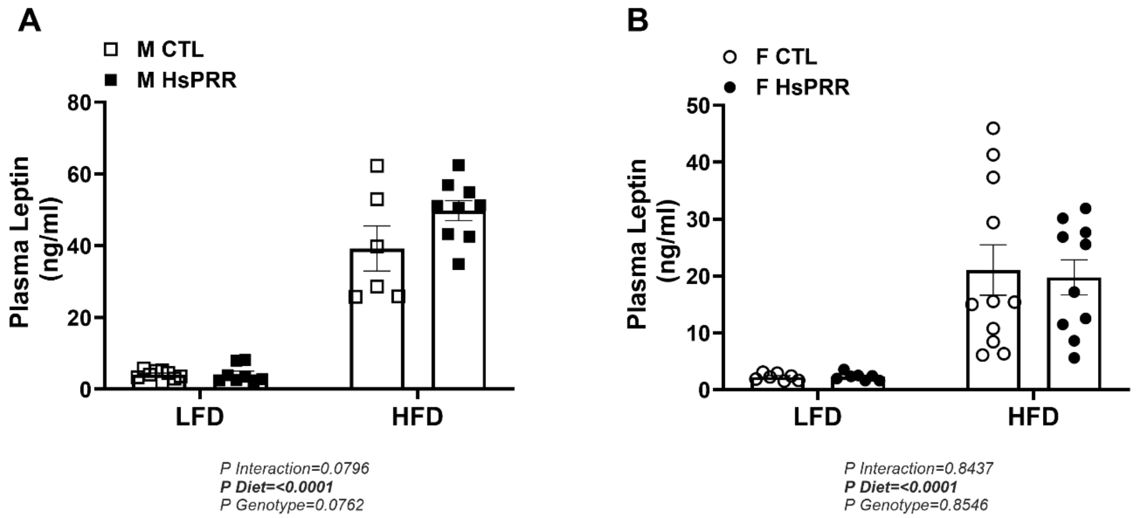

**Supplementary Figure 3 Adi-HsPRRR expression does not affect circulating leptin**

Plasma leptin in (A) Male and (B) Female mice. Data are mean ± SEM of 7-11 mice/group. A 2-way ANOVA was performed to detect differences.

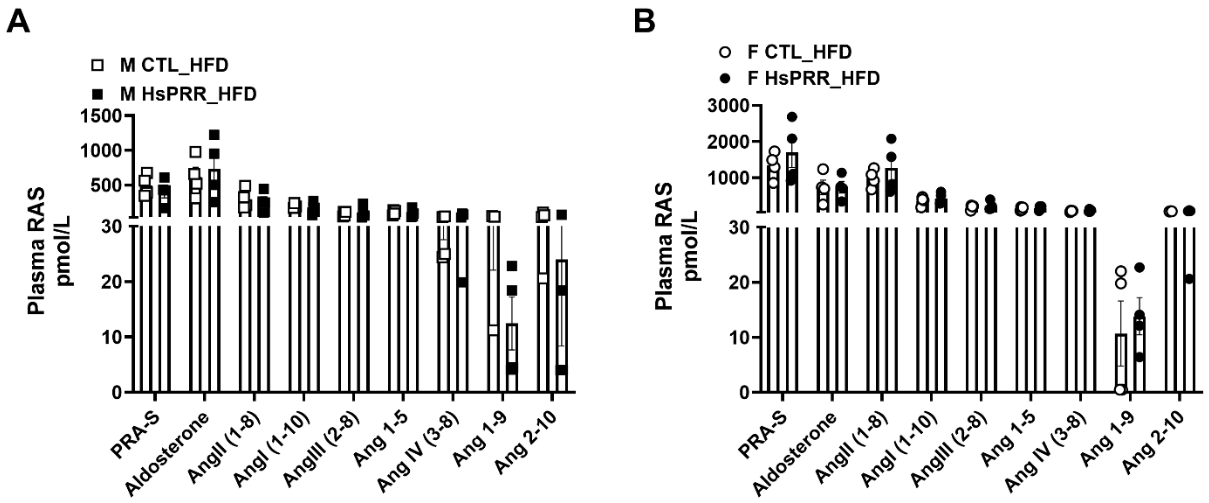


**Supplementary Figure 4 Adi-HsPRRR does not affect other RAS components in obese mice**

Plasma Ang 1-7 in (E) Male and (F) Female mice. Data are expressed as mean ± SEM of 4 mice/group. A t-test was performed to detect differences

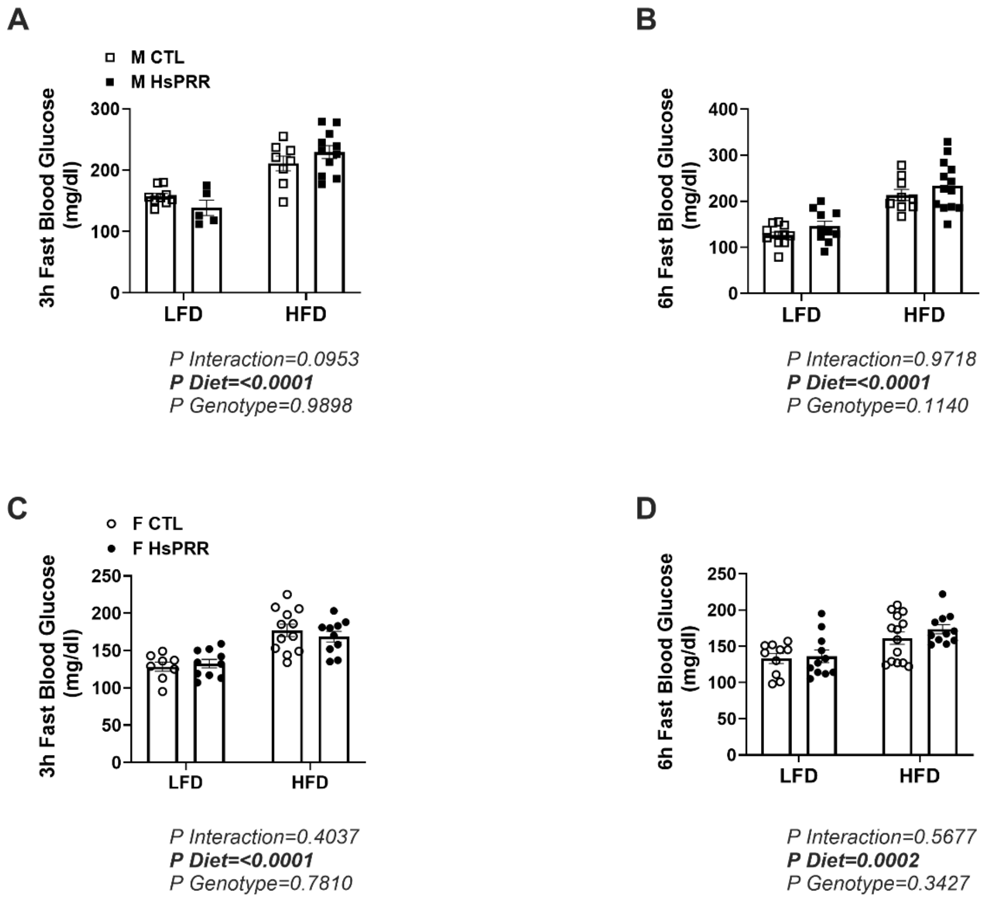


**Supplementary Figure 5 Adi-HsPRRR expression does not affect fasting blood glucose**

Three (3) hours fasting blood glucose in (A) Male and (C) Female mice. Six (6) hours fasting blood glucose in (B) Male and (D) Female mice Data are mean ± SEM of 5-11 mice/group. A 2-way ANOVA was performed to detect differences.

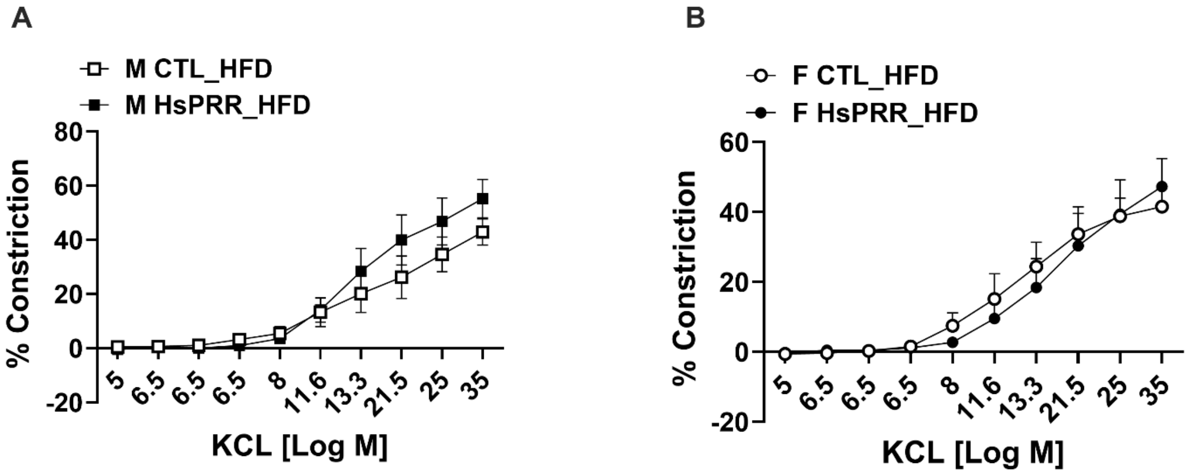


**Supplementary Figure 6 Adi-HsPRR does not change vascular constriction in HFD mice**

Vascular constriction by KCl in (A) Male and (B) Female mice. Data are mean ± SEM of 5 mice/group. A 2-way repeat ANOVA was performed to detect differences.

### Tables

**Supplementary Table 1 Physiological Parameters of Low and High-Fat Diet Mice**

| | LFD CTL | LFD Adi-HsPRR | HFD CTL | HFD Adi-HsPRR |
|---|---|---|---|---|
| MALES | | | | |
| Bodyweight (g) | 28.1±0.6 | 29.9±0.8 | 44.4±2.1 | 46.6±1.6 |
| Fat Mass (%) | 13.1±1.4 | 13.2±1.1 | 40.0±1.2 | 38.0±0.8 |
| Lean Mass (%) | 83.3±1.4 | 82.5±1.4 | 57.7±1.3 | 59.5±0.8 |
| FEMALES | | | | |

| | | | | |
|---|---|---|---|---|
| Bodyweight (g) | 22.8±0.4 | 22.3±0.6 | 38.7±3.1 | 35.6±1.8 |
| Fat Mass (g) | 9.2±0.8 | 9.3±0.5 | 40.5±2.0 | 35.4±2.5 |
| Lean Mass (g) | 85.0±0.7 | 85.7±1.7 | 57.2±2.1 | 60.6±2.7 |

Data are mean ± SEM of 8-16 mice/group; A two-way ANOVA was performed to detect differences. * $P<0.05$ compared with CTL.
